# Detection of melanin-mediated false-positive for BaxΔ2 immunohistochemical staining in human skin tissues

**DOI:** 10.1101/2020.10.30.361956

**Authors:** Sana Basheer, Adriana Mañas, Qi Yao, Katherine Reiner, Jialing Xiang

**Author notes:** Corresponding author: Jialing Xiang, Department of Biology, Illinois Institute of Technology, 3101 South Dearborn Street, Chicago, IL 60616, USA.

## Abstract

BaxΔ2 is a pro-apoptotic isoform of the Bax family. We have previously shown that BaxΔ2 protein expression is low in most healthy human tissues with the protein mainly scattered in the connective tissues. Surprisingly, in skin tissue, the BaxΔ2-positive staining was strikingly strong, especially in the basal layer of the epidermis. It has been documented that melanin in the basal cells displays a brown color that could give false-positive staining in the commonly used DAB-based (3-3’-diaminobenzidine) immunostaining. To avoid the false-positive staining from DAB-melanin, we performed immunofluorescent staining on the same set of tissues which were positive for BaxΔ2 in the DAB immunostaining. The results from co-immunofluorescent staining of BaxΔ2 and a basal cell marker CK-14 showed that the previously detected DAB-stained BaxΔ2-positive epidermal cells were CK-14 positive but BaxΔ2 negative. Consistent with our previous finding, the BaxΔ2-positive cells in the dermis connective tissues are positive in both DAB-based and fluorescence-based immunostainings. These data indicate that the DAB-stained BaxΔ2 positive cells in the basal layer of skin tissue are false-positive due to a misinterpretation of the melanin color, but the dermal connective tissue still presents real BaxΔ2 positive staining.

## Introduction

The skin is the largest organ in the human body, spanning over 22 square feet, and is responsible for about 15-17% of the body mass [1]. It functions in a variety of roles, such as protection from harmful environmental influences and plays an integral part of the immune, endocrine, and nervous systems [2]. Typically, the skin is composed of three distinct layers: the epidermis, the dermis, and the subcutaneous layer. The epidermis consists of 15-20 layers of fully cornified keratinocytes that are constantly renewed as dead skin cells are sloughed away and replenished [3]. The bottom layer of the epidermis, known as the basal layer, consists of epidermal stem cells and melanocytes [4]. The melanocytes are derived from neural crest cells and produce melanin, which can disperse throughout the cells, including the cells of the basal layer [5].

BaxΔ2 is a pro-apoptotic isoform of the Bax family, initially found in patients having colon cancer with high microsatellite instability [6], [7]. Compared to the parental Baxα, BaxΔ2 has a stronger pro-death potency [8]. Unlike most members of the pro-apoptotic Bax family which trigger apoptosis through the mitochondrial pathway, BaxΔ2 proteins are unable to target mitochondria, instead they accumulate in the cytosol as protein aggregates, activating caspase 8-dependent apoptosis [9]–[11]. Expression of BaxΔ2 protein in human tissues has been well characterized by immunohistochemistry with anti-BaxΔ2 antibody [12], [13]. Tissue distribution of BaxΔ2 is also different from that of Baxα: while Baxα is found abundantly and ubiquitously in almost all tissues including normal and cancerous tissue, BaxΔ2 is found in certain types of cells in normal and tumor adjacent normal tissues and rarely detected in malignant tumors [13]. Most BaxΔ2-positive cells in healthy tissues were found scattered in the connective tissues, especially in the colon, ovary, stomach, uterus, breast, and prostate [13].

Expression of BaxΔ2 in skin tissues appeared unique in comparison with other human tissues [13]. In addition to its typical distribution in the dermal connective tissue layer, most of the BaxΔ2-positive cells lined up together at the bottom layer of epidermis [13]. Although the BaxΔ2 antibody has been extensively validated and approved to be specific, we still wondered whether such strong positive staining is real BaxΔ2 [12], [13]. It has been shown that melanin from basal cells is a brown color comparable to the brown oxidized deposits in DAB staining, resulting in a potential false-positive in DAB-based staining [14]. In this study, we analyzed skin tissues by both DAB-immunohistochemistry and immunofluorescent staining. We specifically focused on basal cells using co-immunostaining of both BaxΔ2 and Cytokeratin-14, a basal cell marker, to avoid DAB-mediated artifact during staining.

## 2. Materials and Methods

### 2.1 Materials

Skin FFPE (Formalin-Fixed Paraffin-Embedded) tissue microarray slides were obtained from Biomax (Cat# T211a and SK481) containing both normal and tumor adjacent tissues. The BaxΔ2 monoclonal antibody was generated and validated previously [12], [13]. Anti-Cytokeratin 14 (CK-14) antibody was purchased from Abcam. The secondary antibodies Alexa Fluor 488 and Alexa Fluor 594 were purchased from Life Technologies. Common chemicals mentioned below were obtained from Fischer Scientific. The VECTASTAIN Elite ABC HRP Kit (Peroxidase, Standard) was obtained from Vector Laboratories. The ImmPACT DAB Peroxidase (HRP) Substrate Kit was obtained from Fisher Scientific. The VECTOR Hematoxylin stain was obtained from Vector Labs. The Xylene based mounting media was obtained from Polysciences. Phosphate Buffered Saline (PBS) containing 137 mM NaCl, 2.7 mM KCl, 10 mM Na_2_HPO_4_.7H_2_O, pH 7.3. Blocking buffer was prepared with 3% Bovine Serum Albumin diluted in PBS with 0.1% Tween-20. Nuclear staining basic buffer was prepared with 40 mM NaHCO_3_ and 80 mM MgSO_4_. Fluoromount was obtained from Southern Biotech.

### 2.2 DAB (3,3′-diaminobenzidine) immunohistochemical staining

Tissue sections were deparaffinized in xylene and rehydrated in graded ethanol (100%, 95%, 75%). To remove endogenous peroxidase, tissue slides were incubated in a 3% H_2_O_2_ dilution in PBS at room temperature for 10 minutes. Antigen retrieval was performed by immersing slides in 0.01 M sodium citrate buffer (pH 6.0) at 95°C for 10 minutes. To block non-specific binding, tissue slides were incubated in 3% BSA in PBS containing 0.1% Tween-20. Tissue slides were incubated with primary antibodies overnight at 4°C. The anti-BaxΔ2 antibody was diluted 1:200 in blocking buffer and the anti-CK-14 antibody was diluted 1:300 in blocking buffer. After washing with PBS, slides were incubated in biotin conjugated secondary antibody dilutions of 1:200 in blocking buffer at room temperature for 2 hours. Vectastain ABC Kit (Vector Laboratories) and ImmPACT DAB Peroxidase Substrate Kit (Vector Laboratories) were used to visualize positive-stained cells, while Hematoxylin QS (Vector Laboratories) was used to visualize the nuclei. The tissue slides were then dehydrated in an increasing gradient of ethanol (95%, 100%, 100%) and re-paraffinized with xylene and mounted in Poly-Mount (Polysciences, Inc.). Finally, for image documentation, the slides were scanned using a CRi Pannoramic Scan Whole Slide Scanner with a 40 × NA 0.95 LWD Zeiss objective on high resolution (0.12 μm/pixel) at the Integrated Light Microscopy Core Facility at the University of Chicago and visualized using Pannoramic Viewer 1.15.2.

### 2.3 Immunofluorescent staining

Tissue sections were deparaffinized, rehydrated, antigen retrieved, and blocked in the same way as DAB immunostaining. Tissue slides were incubated with primary antibody dilutions (1:200 for BaxΔ2 and 1:300 for CK-14) overnight at 4°C. After washing with PBS, slides were incubated in Alexa Fluor secondary antibody dilutions (1:200) at room temperature for 2 hours. Tissue slides were stained for nuclei with DAPI (4′,6-diamidino-2-phenylindole) for 10 minutes at room temperature. Slides were incubated in TrueBlack lipofuscin for 10 seconds to quench any autofluorescence. Tissue slides were then mounted with Fluoromount (SouthernBiotech). Immunofluorescent staining was imaged using a BZ-X710 digital microscope (Keyence, Itasca, IL, USA) fitted with 10x, 20x, and 40x (PlanFluor, NA 0.45) objectives.

## 3. Results and Discussion

### 3.1 Expression of BaxΔ2 proteins in normal skin tissues

We have previously shown that expression of BaxΔ2 protein in normal human tissues was very low: the BaxΔ2-positive cells were scattered in the connective tissues in a low rate (1-5%) [13]. However, the skin tissue was an exception. In addition to the detection of BaxΔ2-positive cells in the dermal connective tissue below the epidermis, striking positive staining could be easily visualized in the epidermis [13] (Fig. 1). These positive cells were often clustered in the bottom layer (Fig. 1a) or spread throughout the epidermis (Fig. 1b). We stained 7 normal and 12 tumor adjacent normal tissue samples, and the positive staining was observed in almost all samples examined with different degrees of positive rate and intensity: some were high and strong (Fig. 1a and b) but others were low and light (Fig. 1c). From the tissue distribution and cell morphology, these BaxΔ2-positive stained cells appeared to be basal cells with round nuclei and large cytosol full of positive-stained cytosol granules. These results could imply a unique function of BaxΔ2 in the skin tissue that is different from other tissues. However, we would like to confirm this result before further investigation.

**Fig. 1.**
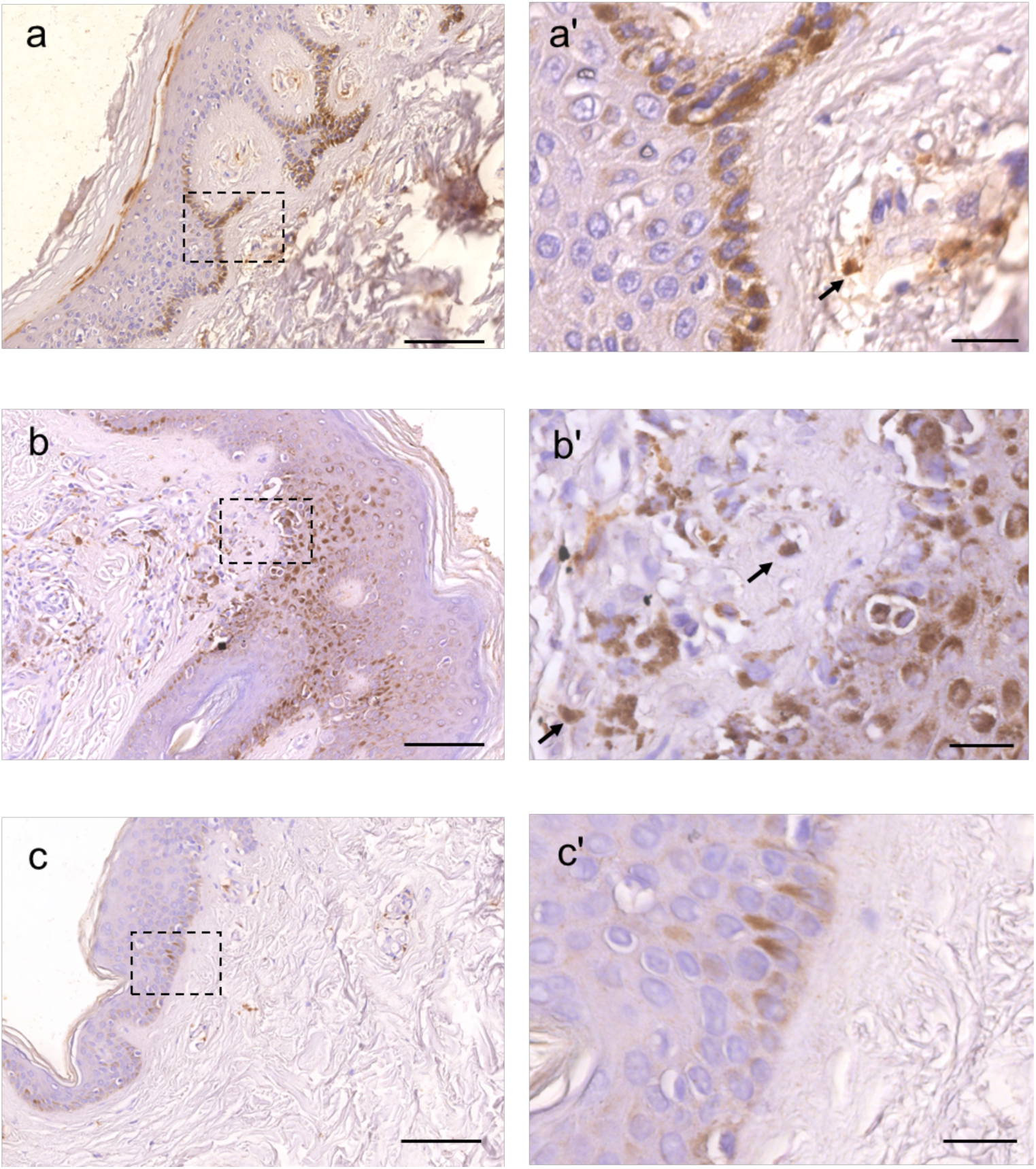
Detection of BaxΔ2 positive staining in normal skin tissues using immunohistochemistry. Human skin tissue slides were stained with anti-BaxΔ2 antibody using standard DAB. Images from three individuals are shown at the left panel (a, b, c). Scale bar, 100 μm. The boxed positive stained areas are enlarged and at the right panel (a’, b’, c’). Black arrows pointed at BaxΔ2 positive cells in dermis connective tissue. Scale bar, 20 μm.

### 3.2 DAB-based BaxΔ2-positive staining in basal cells is false-positive

A survey of literature revealed that excessive amounts of melanin pigments could yield false-positive, as melanin is comparable in color to DAB (3-3’-diaminobenzidine) [14]. Since melanin is produced by the melanocytes and dispersed throughout the basal layer, we wondered whether the positive DAB stained cells were due to melanin rather than BaxΔ2 proteins [15]. To avoid colorimetric based staining, we used a non-colorimetric based staining method: immunofluorescent staining. We co-immunostained the skin tissues for BaxΔ2 and CK-14, which is a hallmark of basal cells in the squamous epithelium [16]. As expected, the CK-14 positive staining showed the typical characteristic pattern in the basal cell layer; the cells were closely stacked and flattened on the bottom layer of the epidermis (Fig. 2a). However, none of the CK-14 positive cells were stained positively for BaxΔ2 (Fig. 2a). BaxΔ2-positive staining was only observed in the dermal connective tissue underneath the basal cell layer (Fig. 2b indicated by arrow), consistent with our previous finding [13].

**Fig. 2.**
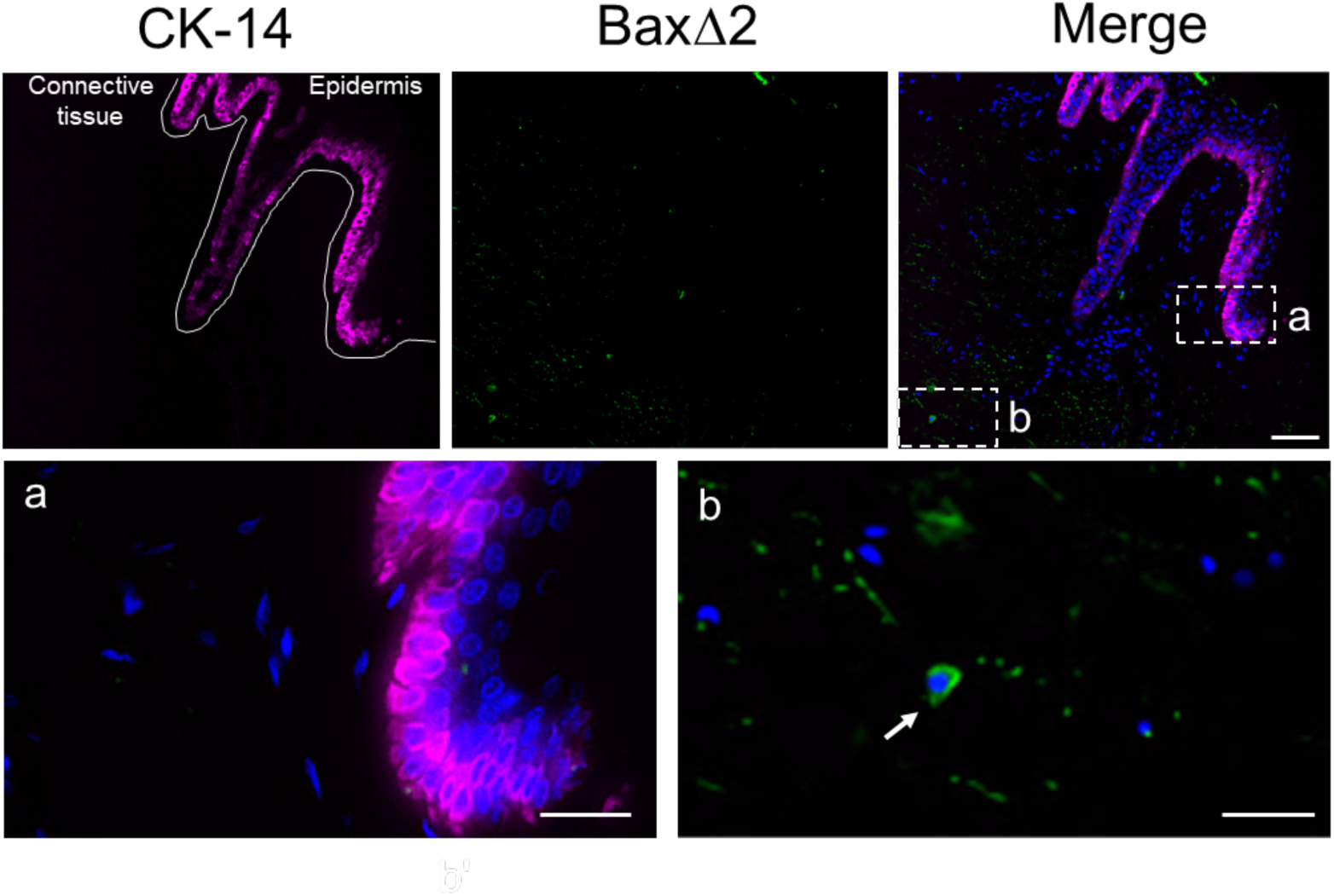
Immunostaining of BaxΔ2 in human skin tissue. Co-immunostaining of normal skin tissues with anti-BaxΔ2 (green) and CK-14 (purple) antibodies. Nuclei were stained with DAPI. The enlarged regions in the boxed areas (a, b) are shown below. Scale bars, 200 μm (top row), 25 μm (bottom row).

These results indicate that the BaxΔ2-positive cells detected in the basal layer by DAB-based immunostaining are false-positives most likely due to a misinterpretation of melanin brown color. Melanin pigments contain sulfur, not iron, making them insoluble in water, and they are found in paraffin embedded tissue even after routine fixation [14]. Our results are consistent with previous reports that the excessive amounts of melanin pigments could hamper histopathologic assessments based on DAB staining [13], [14].

